# Methicillin-Resistant *Staphylococcus aureus* and *bla*_CTX-M_ positive Enterobacteriaceae from retail beef and pork in Singapore

**DOI:** 10.1101/098111

**Authors:** Tse Hsien Koh, Darris Shao Xuan Wong, Nur Khairuddin bin Aron, Quek Choon Lau, Delphine Cao, Li-Yang Hsu

**Affiliations:** Department of Microbiology, Singapore General Hospital, National University of Singapore; School of Life Sciences and Chemical Technology, Ngee Ann Polytechnic, and National University of Singapore; Saw Swee Hock School of Public Health, National University of Singapore

## Abstract

A small study was carried out to see if methicillin-resistant Staphylococcus aureus (MRSA) and Enterobacteriaceae containing *bla*_CTX-M_ extended spectrum beta-lactamase (ESBL) genes could be found in retail meat sold in Singapore. Both were successfully isolated, suggesting the potential for their acquisition in the local community.

---

The spread of antimicrobial resistance is an increasing problem worldwide. The use of antibiotics in farms has resulted in antibiotic resistant bacteria being found in food animals. As Singapore imports most of its food, and therefore has no control of animal husbandry practices, a small study was carried out to see if methicillin-resistant *Staphylococcus aureus* (MRSA) and Enterobacteriaceae containing *bla*_CTX-M_ extended spectrum *β*-lactamase (ESBL) genes could be found in retail meat sold locally.

Beef (n=10) and pork (n=20) were purchased from 15 supermarkets across Singapore. The beef was imported from Australia (n=4), Brazil (n=3), and New Zealand (n=3). The pork was imported from Australia (n=6), Indonesia (n=6), Brazil (n=4), and Malaysia (n=4).

Upon arrival in the laboratory, 25 g of meat was placed in a stomacher bag containing 225 ml of buffered peptone water and macerated in a stomacher for 90 secs at 260 rpm. For MRSA, a 0.6 ml aliquot was added to 3 ml of brain heart infusion broth (BHIB) with 6 mg/L oxacillin and incubated overnight at 35°C. For ESBL-producing Enterobacteriaceae, the remainder in the stomacher bag was incubated overnight at 35°C.

After incubation, 10μl of the incubated broths were diluted 1:100 in buffered peptone water. Then, 10μl of the diluted BHIB was plated onto MRSASelect agar (Bio-Rad) and 10μl of the diluted buffered peptone water from the stomacher bag was plated onto ChromID ESBL (bioMérieux), in duplicate. The plates were incubated at 35°C overnight and later checked for growth. The isolates were identified by MALDI-TOF (Bruker) and susceptibility testing was performed according to CLSI guidelines.^1^ MRSA were typed by multi-locus sequence and spa typing.^2, 3^ CTX-M genes were grouped by multiplex PCR.^4^

We only isolated one MRSA. This was from a pork shoulder imported from Australia. The isolate was resistant to penicillin and cefoxitin, but susceptible to gentamicin, tetracycline, erythromycin, clindamycin, trimethoprim-sulfamethoxazole, rifampicin, fusidic acid, and ciprofloxacin. It was typed to ST3533-MRSA-IV, spa type t015. This novel sequence type is a double locus variant of ST45. The livestock-associated MRSA typically found in pigs are ST398 and ST9.^5^ ST45 is the founder of clonal complex CC45 which is a human clone of MRSA initially described in Europe, but since found worldwide.^5^ This clone is hospital-associated in most countries including Singapore, but may be community-associated in Australia.^6,7^ The fact that this isolate was susceptible to ciprofloxacin is suggestive of community acquisition.

*bla*_ClX-M_-positive Enterobacteriaceae were found in 1 beef and 4 pork samples (Table 1). *bla*_CTX-M-1 group_ is widespread worldwide and appears to be the most common CTX-M gene group found in humans in Singapore and Australia.^8, 9^ The *bla*_CTX-M-8 group_ was first described in Brazil and is relatively uncommon.^10^ We were unable to find any publications reporting *bla*_CTX-M_-positive Enterobacteriaceae isolated from pigs in Australia and Indonesia. However these bacteria (*bla*_CTX-M-1 group_ positive Enterobacteriaceae particularly) have been reported in pigs in other countries.^11^

**Table 1.** Characteristics of ESBL-producing Enterobacteriaceae

| Meat | Strain | Species | Resistance gene | Country of origin | Antimicrobial disk zone diameter (mm)/susceptibility |  |  |  |  |  |  |  |
| --- | --- | --- | --- | --- | --- | --- | --- | --- | --- | --- | --- | --- |
|  |  |  |  |  | ATM | CTX | CTX-CLA | CAZ | CAZ-CLA | CIP | AK | G |
| Beef mince | B3a | *Escherichia coli* | *bla*<sub>CTX-M-1</sub> group | Australia | 18/I | 11/R | 24/na | 21/S | 29/na | 25/S | 22/S | 9/R |
|  | B3b | *Proteus vulgaris* | *bla*<sub>CTX-M-1</sub> group | Australia | 19/I | 13/R | 27/na | 23/S | 30/na | 30/S | 25/S | 10/R |
| Pork mince | P1 | *Escherichia coli* | *bla*<sub>CTX-M-8</sub> group | Australia | 25/S | 20/R | 25/na | 29/S | 32/na | 8/R | 21/S | 22/S |
| Pork shoulder | P2 | *Escherichia coli* | *bla*<sub>CTX-M-1</sub> group | Australia | 16/R | 8/R | 26/na | 17/R | 28/na | 10/R | 25/S | 28/S |
| Pork mince | P6 | *Proteus vulgaris* | *bla*<sub>CTX-M-1</sub> group | Australia | 40/S | 30/S | 35/na | 32/S | 32/na | 35/S | 25/S | 23/S |
| Pork loin | P9 | *Escherichia coli* | *bla*<sub>CTX-M-1</sub> group | Indonesia | 18/I | 10/R | 25/na | 20/I | 30/na | 33/S | 23/S | 25/S |

I, Intermediate; R, Resistant; S, Susceptible; ATM, aztreonam; CTX, cefotaxime; CTX-CLA, cefotaxime-clavulanic acid; CAZ, ceftazidime; CAZ-CLA, ceftazidime-clavulanic acid; CIP, ciprofloxacin; AK, amikacin; G, gentamicin; na, not applicable

This study is limited by a small sample size. Since we did not sample the source animals, we cannot exclude human or environmental contamination occurring during the long journey from farm to retail store. We suspect the MRSA found in pork shoulder represents human contamination. Finding antibiotic resistant bacteria in food products, by itself does not prove that this is the normal route by which humans acquire the resistance determinants. However despite these limitations, we have demonstrated that antibiotic resistance determinants can be found in bacteria isolated from retail meat in Singapore and may potentially result in their acquisition in the community.

## Acknowledgements

We thank Ngee Ann Polytechnic for funding this project.

## References

1. CLSI. Performance Standards for Antimicrobial Susceptibility Testing; 26th informational supplement. CLSI document M100-S26. Clinical and Laboratory Standards Institute, Wayne PA; 2016.

2. Enright MC, Day NPJ, Davies CE, et al. Multilocus sequence typing for characterization of methicillin-resistant and methicillin-susceptible clones of *Staphylococcus aureus*. J Clin Microbiol. 2000; 38: 1008-15.

3. Harmsen D, Claus H, Witte W, et al. Typing of methicillin-resistant *Staphylococcus aureus* in a university hospital setting by using novel software for spa repeat determination and database management. J Clin Microbiol. 2003; 41: 5442–8.

4. Woodford N, Fagan EJ, Ellington MJ. Multiplex pcr for rapid detection of genes encoding CTX-M extended-spectrum *β*-lactamases. J Antimicrob Chemother. 2006; 57:154–5.

5. Bal AM, Coombs GW, Holden MTG, et al. Genomic insights into the emergence and spread of international clones of healthcare-, community- and livestock-associated meticillin-resistant *Staphylococcus aureus:* Blurring of the traditional definitions. J Global Antimicrob Resist. 2016; 6: 95–101.

6. Hsu LY, Harris SR, Chlebowicz MA, et al. Evolutionary dynamics of methicillin-resistant *Staphylococcus aureus* within a healthcare system. Genome Biol. 2015; 53:146–9.

7. Coombs GW, Daley DA, Lee YT, et al. Australian group on antimicrobial resistance Australian *Staphylococcus aureus* sepsis outcome programme annual report, 2014. Commun Dis Intell. 2016; 40:E244–54.

8. Tan TY, Ng LSY, He J, et al. CTX-M and ampC *β*-lactamases contributing to increased prevalence of ceftriaxone-resistant *Escherichia coli* in Changi General Hospital, Singapore. Diagn Microbiol Infect Dis. 2010; 66: 210–3.

9. Zong Z, Partridge SR, Thomas L, et al. Dominance of blaCTX-M within an Australian extended-spectrum *β*-lactamase gene pool. Antimicrob Agents Chemother. 2008; 52: 4198–202.

10. Bonnet et al. A novel CTX-M *β*-lactamase (CTX-M-8) in cefotaxime-resistant Enterobacteriaceae isolated in Brazil. Antimicrob Agents Chemother. 2000; 44:1936–42.

11. Seiffert SN, Hilty M, Perreten V, Endimiani A. Extended-spectrum cephalosporin-resistant gram-negative organisms in livestock: An emerging problem for human health? Drug Resist Updat. 2013; 16: 22–45.

